# Exploring Functional Brain Network Modularity in Educational Contexts

**DOI:** 10.1101/2022.01.06.475275

**Authors:** Adam B. Weinberger, Robert A. Cortes, Richard F. Betzel, Adam E. Green

## Abstract

The brain’s modular functional organization facilitates adaptability. Modularity has been linked with a wide range of cognitive abilities such as intelligence, memory, and learning. However, much of this work has (1) considered modularity while a participant is at rest rather than during tasks conditions and/or (2) relied primarily on lab-based cognitive assessments. Thus, the extent to which modularity can provide information about real-word behavior remains largely unknown. Here, we investigated whether functional modularity during resting-state and task-based fMRI was associated with academic learning (measured by GPA) and ability (measured by PSAT) in a large sample of high school students. Additional questions concerned the extent to which modularity differs between rest and task conditions, and across spatial scales. Results indicated that whole-brain modularity during task conditions was significantly associated with academic learning. In contrast to prior work, no such associations were observed for resting-state modularity. We further showed that differences in modularity between task conditions and resting-state varied across spatial scales. Taken together, the present findings inform how functional brain network modularity – during task conditions and while at rest – relate to a range of cognitive abilities.

## Introduction

The brain is able to adaptively reconfigure its functional connections to meet the demands of daily life and perform a wide variety of cognitive tasks. A growing body of evidence indicates that such adaptability stems from the brain’s modular organization (Bassett & Mattar, 2017; Betzel, Gu, et al., 2016; Medaglia et al., 2015; Power et al., 2011; Sporns, 2010; Sporns & Betzel, 2016; Yeo et al., 2011; Toga et al., 2006). Modularity, a measure of network segregation, has been proposed as a basis of cognitive abilities such as general or fluid intelligence (Barbey, 2018; Chiappe & MacDonald, 2005; Finn et al., 2015; Hilger et al., 2020, 2020), memory (Stanley et al., 2014; Stevens et al., 2012; Wig, 2017), and learning (Arnemann et al., 2015; Baniqued et al., 2018; Bassett & Mattar, 2017; Duncan & Small, 2016; Gallen et al., 2016; Gallen & D’Esposito, 2019). Given these associations, a number of recent perspectives suggest that modularity may have utility as a neural biomarker (Barbey, 2018; Bassett & Mattar, 2017; Gabrieli et al., 2015; Gallen & D’Esposito, 2019; Hilger et al., 2017, 2020). However, a priority for cognitive neuroscience is to better establish how modularity can be utilized in such a manner. For example, what is the correspondence between an individual’s intrinsic modularity (i.e., at rest or in the absence of external demands) and their modular organization during task conditions? Is task-based modularity a better predictor of cognitive ability than is modular organization at rest? Can modularity reliably predict cognitive performance across contexts? The present study sought to address these outstanding questions.

A modular network is one in which network “nodes” are clustered into multiple distinct “subgraphs” or “communities”, with dense connectivity among nodes within each subgraph and comparatively sparser connections across subgraphs (Newman, 2006; Sporns & Betzel, 2016). In other words, nodes within a given subgraph communicate with one another more than they do with nodes of other subgraphs. Modularity, then, is a quantification of the extent to which a network can be divided into distinct subgraphs. The brain is one such modular network, with nodes typically referring to specific brain regions and connectivity derived from structural tracts (i.e., white matter; “structural modularity”) or functional associations (i.e., functional connectivity; “functional modularity”). The brain’s modular organization is found across all spatial scales; discrete modules could be conceptualized as the two cerebral hemispheres, coordinated networks (e.g., frontoparietal), smaller networks (frontal, parietal), gray matter nuclei, and cell columns (Sporns, 2010).

Modularity has been identified in species ranging from the nematode *C. elegan* (Pan et al., 2010; Sohn et al., 2011) to more complex organisms and systems (Betzel et al., 2017; Harriger et al., 2012; Hilgetag & Young, 2000; Modha & Singh, 2010; Sporns & Betzel, 2016). The prevalence of modularity across scales and lifeforms suggests that such organizational properties facilitate evolutionary fitness (Bassett & Bullmore, 2006; Betzel et al., 2017; Gerhart & Kirschner, 2007; Kirschner & Gerhart, 1998; Sporns & Betzel, 2016). According to these perspectives, modularity increases the brain’s capacity for independent rearrangement of subgraphs in response to external experiences (Kashtan et al., 2007; Kashtan & Alon, 2005), which, in turn, enables greater system-wide adaptability (Bassett & Bullmore, 2006; Mattar et al., 2018; Meunier et al., 2009). Thus, modularity provides robustness (because of reduced constraints on any single module; Mattar et al., 2018) as well as “evolvability” (i.e., the capacity to adaptively generate variation in organization; Anderson & Finlay, 2014). The result, then, is that neuronal populations that are not specialized to respond to a given input can remain unchanged while task-relevant networks reconfigure and generate appropriate outputs.

Given the above, modularity at rest should differ from modularity when an individual is engaged in an external task. More specifically, because task demands elicit reconfiguration of functional connectivity between relevant subgraphs that may be otherwise segregated (e.g., Cole et al., 2014; Medaglia et al., 2015; Power et al., 2011; Yeo et al., 2011), modularity should decrease for task conditions (i.e., relative to rest or intrinsic organization). That is, subgraphs should be more segregated in the absence of external demands. A growing body of literature indicates that this likely to be the case (Cohen & D’Esposito, 2016; Di et al., 2013; Finc et al., 2017, 2020; Kitzbichler et al., 2011). For example, Finc and colleagues (2020) found that modularity was higher at rest than during an n-back task and parametrically decreased with greater working memory load, suggesting that the brain forms functional connections between task-relevant modules that may be less likely to communicate at rest. The finding that modularity was inversely related to working memory load further indicates that more challenging conditions may bring about even greater departure from intrinsic organization. This interpretation accords with the Global Workspace Theory (Dehaene et al., 1998), which argues for a distinction between an intrinsic “global” workspace and specialized, integrated modules for specific tasks. This line of thinking suggests that less challenging tasks can be successfully performed within segregated modules (i.e., preserving intrinsic modularity) compared to more challenging tasks that necessitate greater integration between more disparate subgraphs (i.e., resulting in lower brain-wide modularity). In the present study, we further investigated task-based changes in modularity by comparing modular organization at rest with modular organization during two cognitive tasks.

Other recent work indicates that modularity facilitates learning (Bassett & Mattar, 2017), consistent with the perspective that modularity promotes adaptability. In artificial intelligence, modularity has been shown to support task learning (Jo et al., 2018) and the acquisition of new skills without catastrophic forgetting, the sudden loss of previously acquired information (Ellefsen et al., 2015; McCloskey & Cohen, 1989). Similar findings have been observed in humans. In one such study, researchers investigated whether resting-state modularity was predictive of improved functioning in patients with acquired brain injury following a five-week cognitive training program (Arnemann et al., 2015). Results indicated a significant, positive association between modularity and improvement in executive function and attention, suggesting modularity may be a marker of intervention-related plasticity. Similarly, in a study of older adults, people with more modular brains (again, at rest) showed greater cognitive improvements after a 12-week intervention targeting attention, reasoning, and innovation (Gallen et al., 2016). Resting-state modularity has also been correlated with executive function gains following an exercise-based clinical trial (Baniqued et al., 2018) and working memory training (Iordan et al., 2018) in older adults, as well as the effectiveness of imitation-based therapy for aphasic patients (Duncan & Small, 2016). More recent studies have begun to establish a similar pattern of results in younger, healthier samples. For example, children with higher resting-state modularity exhibited greater improvements across a range of cognitive tests following an intervention to improve working memory and reasoning (Baniqued et al., 2019). In another study of young children, resting-state modularity was positively associated with improved cognitive performance after participation in an extended physical fitness program (Chaddock-Heyman et al., 2020).

Although the literatures on modularity and learning are broadly cohesive, they are also somewhat limited in scope. First, prior work examining modularity and learning has relied heavily on resting-state fMRI, despite differences in modular organization between task and rest conditions^1^. Since challenging tasks are typically accompanied by a reduction in modularity, it is plausible that individual differences in the extent to which one can maintain higher modularity *despite task demands* may be even more reflective of cognitive abilities. That is, higher *task-based* modularity may signal less of a need to deviate from one’s intrinsic workspace (Dehaene et al., 1998). The present study investigated this possibility by relating learning to both resting-state and task-based modularity. More broadly, the inclusion of resting-state and task-based fMRI aligns with a growing emphasis placed on “multimodal” protocols to better understand how principles of brain organization relate to cognitive performance (e.g., Baykara et al., 2021).

A second limitation of prior work relating modularity to learning is that research has been largely restricted to patients with brain injury and older individuals (see Baniqued et al., 2019; Chaddock-Heyman et al., 2020 as recent exceptions). These populations are likely to have impaired cognitive functioning and reduced modularity (Betzel et al., 2014; Geerligs et al., 2015; Song et al., 2014), particularly for brain systems involved in executive functioning and associative processing (e.g., frontoparietal control network, ventral attention network; Chan et al., 2014). It is plausible, then, that individual differences in modularity may be a downstream result of – or at least meaningfully influenced by – neurological damage or degeneration. That is, modularity may be a proxy for “brain health” rather than uniquely related to a larger organizational framework, and those with less significant decay would show higher modularity and be more receptive to interventions. This possibility is supported by work showing a correlation between modularity and learning for older, but not younger adults (Duncan & Small, 2016).

The existing literature is also limited in the kind of learning that has been investigated. For instance, despite the value placed on classroom-based education and its connection to achievement later in life (French et al., 2015; Noble & Sawyer, 2004; Noble & Sawyer, 2002; Sawyer, 2013; Schulenberg et al., 1994) – little is presently known concerning the relationship between modularity and in-school learning/achievement. Identifying the neurocognitive processes supportive of school-based learning has the potential to yield significant real-world value.

Here, we applied network analyses of fMRI data to examine modularity at rest and during two cognitive tasks in a group of high school students. Key questions concerned: (i) how resting-state modularity compared with task-based modularity; (ii) whether task-based modularity was associated with task performance; and (iii) if modularity – either at rest or during either task – was associated with academic learning and ability.

## Methods

### Participants

Seventy-nine students (M_age_ = 16.70 years; SD = 0.49 years, 50.63% female, 49.37% male) were recruited from five Northern Virginia public high schools prior to the start of their senior year. All participants were right-handed and reported no history of mental illness or psychoactive drug use. Written informed consent was obtained from each participant as well as assent from a parent or guardian. All procedures received approval from the Georgetown University IRB.

### Experimental procedures

The study was performed at the Center for Molecular Imaging at Georgetown University. Participants performed a verbal relational reasoning task (henceforth “reasoning”) and a mental rotation task (MRT; order counterbalanced) inside the MRI scanner. Tasks were selected based on separate hypotheses described elsewhere (Cortes et al., 2021). The imaging sessions also included a 5-minute resting-state scan as well as a high-resolution anatomical image acquisition (MPRAGE). We did not obtain valid imaging data for all tasks for all participants due to factors such as subject movement (see Additional Processing for Network Construction) and time constraints at the time of data collection. In order to increase statistical power, we elected to use all available data for each participant rather than restrict analyses to only the full set of participants with complete data for all three fMRI conditions (i.e., resting state, reasoning, MRT), as this would have substantially reduced the sample size. The total N for each measure is detailed in Table S1.

### Verbal relational reasoning task

Participants completed 60 verbal relational reasoning problems (“reasoning”) in the fMRI scanner. Each problem was composed of two premises and one conclusion. Instructions stated that participants should judge whether the information in the conclusion necessarily followed from the premises or not (e.g., Premise 1: “The dog is smarter than the cat”; Premise 2: “The cat is smarter than the bird”; Conclusion: “The dog is smarter than the bird”; Fig. S1). Participants also completed 20 trials of a control task in which they determined if the conclusion exactly matched one of the premises. Participants had 8 seconds to respond after the presentation of the conclusion. Task performance was estimated using a composite measure of speed and accuracy (see SI for calcuation of this variable; Vandierendonck, 2017; Woltz & Was, 2006).

### Mental rotation task

Participants completed a version of the Mental Rotation Task (MRT; Peters & Battista, 2008; Shepard & Metzler, 1971) for which they assessed whether two images of 3-dimensional objects (Fig. S1) were rotated images of the same object or images of different objects. Objects were shown at three different angles (50, 100, 150 degrees) to vary task difficulty. We also included 12 trials with 0 degrees of rotation as a control condition. There were 84 trials in total. Participants had 7 seconds to respond. Performance was captured using a composite speed-accuracy metric.

### Markers of academic ability and learning

Finally, we obtained participants’ Grade Point Average (GPA) and Preliminary Scholastic Aptitude Test (PSAT) scores prior to the scanning session. PSAT scores are used to predict performance at the collegiate level and, in the present study, were conceptualized a correlate of academic ability and fluid intelligence (Coyle, 2015; Lent et al., 1986). Student GPA was considered a proxy for classroom learning (Duff et al., 2004; Komarraju et al., 2011).

### fMRI image acquisition and processing

#### Imaging procedures

Image acquisition was performed on a 3 T Siemens Trio Tim MRI scanner. Task fMRI data were acquired from T2*-weighted echoplanar imaging (EPI) sequence (37 3.0 mm transversal slices; 64 x 64 matrix; repetition time = 2000 ms; echo time = 30 ms; field of view = 192 mm; 3.0 x 3.0 x 3.0 mm voxels; flip angle = 90 degrees). To account for magnet stabilization, the first 2 volumes were excluded from analysis. Task were presented with E-Prime and viewed through a mirror attached to a head coil. Participants used SR-boxes in their right hand (index and middle finger) to respond. Resting-state fMRI data (5 minutes in total) were collected using a T2*-weighted EPI sequence (43 2.55 mm transversal slices; 64 x 64 matrix; repetition time = 2280 ms; echo time = 30 ms; field of view = 192 mm; 3.0 x 3.0 x 2.5 mm voxels; flip angle = 90 degrees). A high-resolution T1-weighted MPRAGE image (176 1.00 mm slices; 256 x 256 matrix; repetition time = 1900 ms; echo time = 2.52 ms; field of view = 250 mm; 1.0 x 1.0 x 1.0 mm; flip angle = 9 degrees) was obtained for registration of functional data.

### fMRI data preprocessing

Results included in this manuscript come from preprocessing performed using fMRIPrep 1.5.0 (Esteban et al., 2017, 2019; RRID:SCR_016216), which is based on Nipype 1.2.2 (Gorgolewski et al., 2011; Gorgolewski et al., 2018; RRID:SCR_002502)

### Anatomical data preprocessing

The T1-weighted (T1w) image was corrected for intensity non-uniformity (INU) with N4BiasFieldCorrection (Tustison et al., 2010), distributed with ANTs 2.2.0 (Avants et al., 2008, RRID:SCR_004757), and used as T1w-reference throughout the workflow. The T1w-reference was then skull-stripped with a Nipype implementation of the antsBrainExtraction.sh workflow (from ANTs), using OASIS30ANTs as target template. Brain tissue segmentation of cerebrospinal fluid (CSF), white-matter (WM) and gray-matter (GM) was performed on the brain-extracted T1w using fast (FSL 5.0.9, RRID:SCR_002823, Zhang et al., 2001). Volume-based spatial normalization to two standard spaces (MNI152NLin6Asym, MNI152NLin2009cAsym) was performed through nonlinear registration with antsRegistration (ANTs 2.2.0), using brain-extracted versions of both T1w reference and the T1w template. The following templates were selected for spatial normalization: FSL’s MNI ICBM 152 non-linear 6th Generation Asymmetric Average Brain Stereotaxic Registration Model (Evans et al., 2012, RRID:SCR_002823; TemplateFlow ID: MNI152NLin6Asym), ICBM 152 Nonlinear Asymmetrical template version 2009c (Fonov et al., 2009, RRID:SCR_008796; TemplateFlow ID: MNI152NLin2009cAsym).

### Functional data preprocessing

For each of the BOLD runs found per subject (reasoning, MRT, and rest), the following preprocessing was performed. First, a reference volume and its skull-stripped version were generated using a custom methodology of fMRIPrep. The BOLD reference was then co-registered to the T1w reference using flirt (FSL 5.0.9, Jenkinson & Smith, 2001) with the boundary-based registration (Greve & Fischl, 2009) cost-function. Co-registration was configured with nine degrees of freedom to account for distortions remaining in the BOLD reference. Head-motion parameters with respect to the BOLD reference (transformation matrices, and six corresponding rotation and translation parameters) are estimated before any spatiotemporal filtering using mcflirt (FSL 5.0.9, Jenkinson et al., 2002). BOLD runs were slice-time corrected using 3dTshift from AFNI 20160207 (Cox & Hyde, 1997, RRID:SCR_005927). The BOLD time-series (including slice-timing correction when applied) were resampled onto their original, native space by applying a single, composite transform to correct for head-motion and susceptibility distortions. These resampled BOLD time-series will be referred to as preprocessed BOLD in original space, or just preprocessed BOLD. The BOLD time-series were resampled into several standard spaces, correspondingly generating the following spatially-normalized, preprocessed BOLD runs: MNI152NLin6Asym, MNI152NLin2009cAsym. First, a reference volume and its skull-stripped version were generated using a custom methodology of fMRIPrep. Automatic removal of motion artifacts using independent component analysis (ICA-AROMA, Pruim et al., 2015) was performed on the preprocessed BOLD on MNI space time-series after removal of non-steady state volumes and spatial smoothing with an isotropic, Gaussian kernel of 6mm FWHM (full-width half-maximum). Corresponding “non-aggressively” denoised runs were produced after such smoothing. Additionally, the “aggressive” noise-regressors were collected and placed in the corresponding confounds file. Several confounding time-series were calculated based on the preprocessed BOLD: framewise displacement (FD), DVARS and three region-wise global signals. FD and DVARS are calculated for each functional run, both using their implementations in Nipype (following the definitions by Power et al., 2014). The three global signals are extracted within the CSF, the WM, and the whole-brain masks. Additionally, a set of physiological regressors were extracted to allow for component-based noise correction (CompCor, Behzadi et al., 2007). Principal components are estimated after high-pass filtering the preprocessed BOLD time-series (using a discrete cosine filter with 128s cut-off) for the two CompCor variants: temporal (tCompCor) and anatomical (aCompCor). tCompCor components are then calculated from the top 5% variable voxels within a mask covering the subcortical regions. This subcortical mask is obtained by heavily eroding the brain mask, which ensures it does not include cortical GM regions. For aCompCor, components are calculated within the intersection of the aforementioned mask and the union of CSF and WM masks calculated in T1w space, after their projection to the native space of each functional run (using the inverse BOLD-to-T1w transformation). Components are also calculated separately within the WM and CSF masks. For each CompCor decomposition, the k components with the largest singular values are retained, such that the retained components’ time series are sufficient to explain 50 percent of variance across the nuisance mask (CSF, WM, combined, or temporal). The remaining components are dropped from consideration. The head-motion estimates calculated in the correction step were also placed within the corresponding confounds file. The confound time series derived from head motion estimates and global signals were expanded with the inclusion of temporal derivatives and quadratic terms for each (Satterthwaite et al., 2013). Frames that exceeded a threshold of 0.5 mm FD or 1.5 standardised DVARS were annotated as motion outliers. All resamplings can be performed with a single interpolation step by composing all the pertinent transformations (i.e. head-motion transform matrices, susceptibility distortion correction when available, and co-registrations to anatomical and output spaces). Gridded (volumetric) resamplings were performed using antsApplyTransforms (ANTs), configured with Lanczos interpolation to minimize the smoothing effects of other kernels (Lanczos, 1964). Non-gridded (surface) resamplings were performed using mri_vol2surf (FreeSurfer).

Many internal operations of fMRIPrep use Nilearn 0.5.2 (Abraham et al., 2014, RRID:SCR_001362), mostly within the functional processing workflow. For more details of the pipeline, see the section corresponding to workflows in fMRIPrep’s documentation.

### Additional processing for network construction

Following fMRIPrep image preprocessing, additional processing of individual subject functional data was conducted with XCP Engine (Ciric et al., 2018). Processing was performed using ICA-AROMA (Pruim et al., 2015) with global signal regression. ICA-AROMA identifies sources of variance in brain signal using independent component analyses, incorporating each source as either noise or signal of interest. Sources identified as noise are then denoised along with mean signal from white matter, CSF, and grey matter (because of global signal regression). This processing pipeline was chosen in order to aggressively minimize the influence of head motion (Macey et al., 2004; Pruim et al., 2015), which has been shown to have particularly profound effects on functional connectivity analyses (Power et al., 2012; Van Dijk et al., 2012). Further, each volume was motion censored, and volumes with >0.5mm of motion were flagged. Participants with > 25% bad volumes were excluded from analyses. Frame-to-frame displacement (i.e., mean framewise displacement; FD) for all participants was retained for use in subsequent linear regression models. Additional processing steps included demeaning and detrending, mathematical expansion to model delayed or nonlinear signal fluctuations, temporal filtering [0.01 - 0.1 Hz], and spatial smoothing. Finally, we regressed stimulus/task-evoked activity during both tasks to study residual, task-free/“state” signal (Cole et al., 2014; Gratton et al., 2016; Hwang et al., 2019; Norman-Haignere et al., 2012).

### Atlas/ROI selection

We segmented the brain into regions of interest (ROIs) using two standardized atlases. The cortical surface was divided in 200 parcellations based on functional homogeneity (Schaefer et al., 2018). Subcortical ROIs – included because of their well-established role in reasoning (Christoff et al., 2001; Melrose et al., 2007) and spatial thinking (Gao et al., 2017; Packard & Knowlton, 2002; Sukumar et al., 2012) – were obtained from the anatomically-defined Harvard-Oxford atlas (Desikan et al., 2006). While the methods of parcellation were not consistent across these two atlases (i.e., functionally-based for cortical; anatomically-based for subcortical), we did not have any *a priori* hypotheses regarding the involvement of specific regions of subcortical structures, and therefore preferred to treat each structure as a discrete ROI. Further, we used a relatively coarse cortical atlas to partially mitigate differences in parcel size between cortical and subcortical ROIs.

### Network analyses

#### Functional connectivity

Functional connectivity was estimated using the signal timeseries of the 216 ROIs (200 cortical parcels from Schaefer et al., 2018; 16 subcortical structures from Harvard-Oxford Atlas; Desikan et al., 2006). For each participant, we performed pairwise correlations for every ROI to every other ROI, resulting in a 216 x 216 (z-transformed), weighted functional connectivity matrix. Separate functional connectivity matrices were obtained for reasoning, MRT, and rest.

#### Modularity

Modularity is characterized by a clustering of nodes (i.e., ROIs) into multiple and distinct subgraphs or communities, with a greater number of connections among nodes with a community than connections between communities (Genon, Reid, Langner, Amunts, & Eickhoff, 2018; Lorenz et al., 2017; Power et al., 2011; Toga, Thompson, Mori, Amunts, & Zilles, 2006; Yeo et al., 2011). In order to obtain an estimate of functional modularity, we used an increasingly prevalent method known as modularity maximization (Newman & Girvan, 2004), which assigns nodes to communities by comparing the connection density for each node (i.e., the portion of possible edges that are actual connections) against the connection density for each node of a generated null model. Modularity maximization algorithms then attempt to place nodes connected by edge weights that exceed what is found in the null model into the same community, such that the internal connection density will maximally exceed the internal density of the null model. This ultimately produces a modularity statistic (“Q”), which measures the extent to which communities or subgraphs exhibit segmentations/clustering (higher Q = higher modularity; Betzel & Bassett, 2016; Newman, 2006). Because the algorithm is non-deterministic and can only approximate maximal modularity, we repeated the process 100 times per participant per imaging task to increase statistical robustness, and used consensus clustering to create a consensus community partition. Modularity was thus calculated as average Q across the 100 iterations.

While some prior work has relied on binary representations of functional connectivity (i.e., nodes that are correlated above some threshold are said to be connected, those below it are not), binarizing functional connections may introduce a number of statistical complications, such as the need to create matrices over multiple thresholds as well as the loss of potentially meaningful negative correlations (Rubinov & Sporns, 2011). In order to avoid these consequences and more fully characterize functional networks, we used the original weighted functional connectivity matrices to obtain modularity estimations. Consistent with recommended procedures (Rubinov & Sporns, 2011), Q was calculated using a negative asymmetrical weight that treats negative correlations as auxiliary to positive correlations, as the latter more directly relate to modular organization (i.e., positive correlations among nodes directly indicate that those nodes correspond to a module whereas negative correlations can only dissociate nodes from modules). We therefore defined modularity (here, Q^*^) as

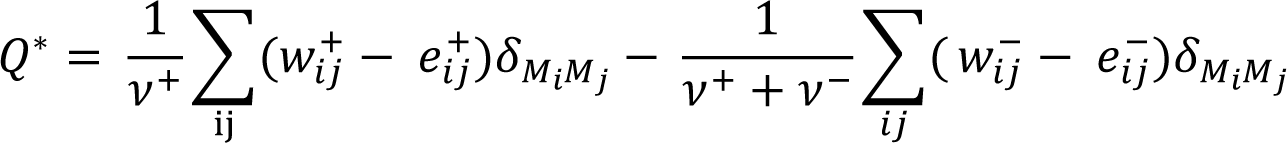

where the left side of the equation shows the calculation of modularity for positive connections only 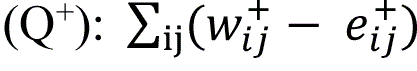 is the average difference between observed within-module positive connection weights, 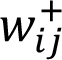, and those expected by chance. 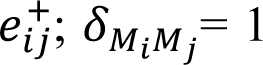 when nodes *i* and *j* are in the same module and 0 when they are not; the positive-weight scaling factor, 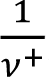, adjusts Q^+^ to fall between 0 and 1. The right side of the equation reflects calculation of Q^-^, modularity based on negative weights, rescaled by the total connection weight, 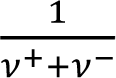. Note that the difference in the scaling factor for Q^+^ and Q^-^ is used to place less weight on the role of negative connections in the calculation of Q^*^.

Further, it is evident from the above calculation that modularity is heavily influenced by “lower level” network features. That is, the number (and average weight) of positive and negative edges strongly influence the overall calculation of Q. In order to better understand the influence of these network characteristics, we retained the number and average weight of positive and negative edges for each participant. These values were plotted against Q (Table S2), and were considered for use as additional covariates in subsequent linear regression models.

Crucially, modularity exists over multiple scales, meaning that within any one community there are a number of smaller, segregated communities, and that these communities within communities also display modular organization. One notable shortcoming to modularity maximization stems from what is known as the resolution limit – an insensitivity to this inherently hierarchical organization. Theoretically, one could choose to “cut” at any point and observe the extent of segmentation at that point. The resolution parameter, γ (Reichardt & Bornholdt, 2006), can be used to mitigate this issue. By changing γ, one is able to vary the threshold for segmenting the brain into communities of different sizes (Betzel & Bassett, 2016). When γ < 1, modularity maximization detects larger (and fewer) communities, resulting in a higher value for Q; when γ > 1 the algorithm produces several smaller communities and a smaller Q statistic (Fig. 1). Although we did not have an *a priori* reason to focus on modularity at any particular scale in the present study, we calculated Q across a range of γ – from 0.5 to 2.0 in increments of 0.5 – to provide a more complete picture of participant modularity. Modularity calculations were performed using functions in Brain Connectivity Toolbox (https://sites.google.com/site/bctnet/; Rubinov & Sporns, 2010). All custom MATLAB code used for analyses supporting the results of the present study are available on the Open Science Framework (https://osf.io/naj3y/).

**Figure 1.**
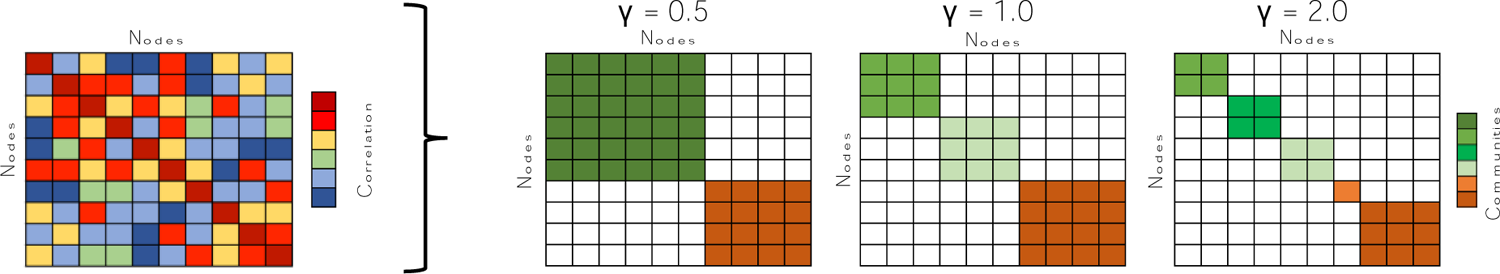
Toy graph of modularity over multiple scales. A functional connectivity matrix (left) is used to obtain a modularity statistic (Q). Using the resolution parameter, γ, modularity is calculated over varying scales (right). As γ increases, the number of communities increase while the size of the communities decreases.

### Results Modularity during rest and task conditions

We used the technique of modularity maximization to estimate modular organization at rest and during two task conditions (reasoning, MRT) across four resolution parameters (γ from 0.5 to 2.0, at 0.5 intervals; Fig. 1). As γ increases, Q is expected to decrease because the modularity maximization algorithm is biased towards more (and smaller) communities.

We first investigated whether the task conditions altered modularity by comparing Q_Rest_ with Q_Reasoning_ and Q_MRT_. Across all resolution parameters, paired t-tests revealed that modularity was significantly higher at rest than during reasoning (all *t*(63) > 9.60, *p* < 0.0001; Fig. 2A), with the nominally greatest difference at γ = 0.5 (Q_Rest_ = 0.57, *SD* = 0.04; Q_Reasoning_ = 0.49, *SD* = 0.05). This result suggests that task demands during reasoning caused a reconfiguration of – and decrease in – intrinsic (i.e., resting-state) modular organization. Q_Rest_ was also significantly greater than Q_MRT_ for γ = 0.5 (*t*(69) = 3.71, *p* = 0.004) and γ= 1.0 (*t*(69) = 2.55, *p* = 0.01), but not for γ = 1.5 (*t*(69) = 1.73, *p* = 0.09) or γ = 2.0 (*t*(69) = 1.43, *p* = 0.16).

**Figure 2.**
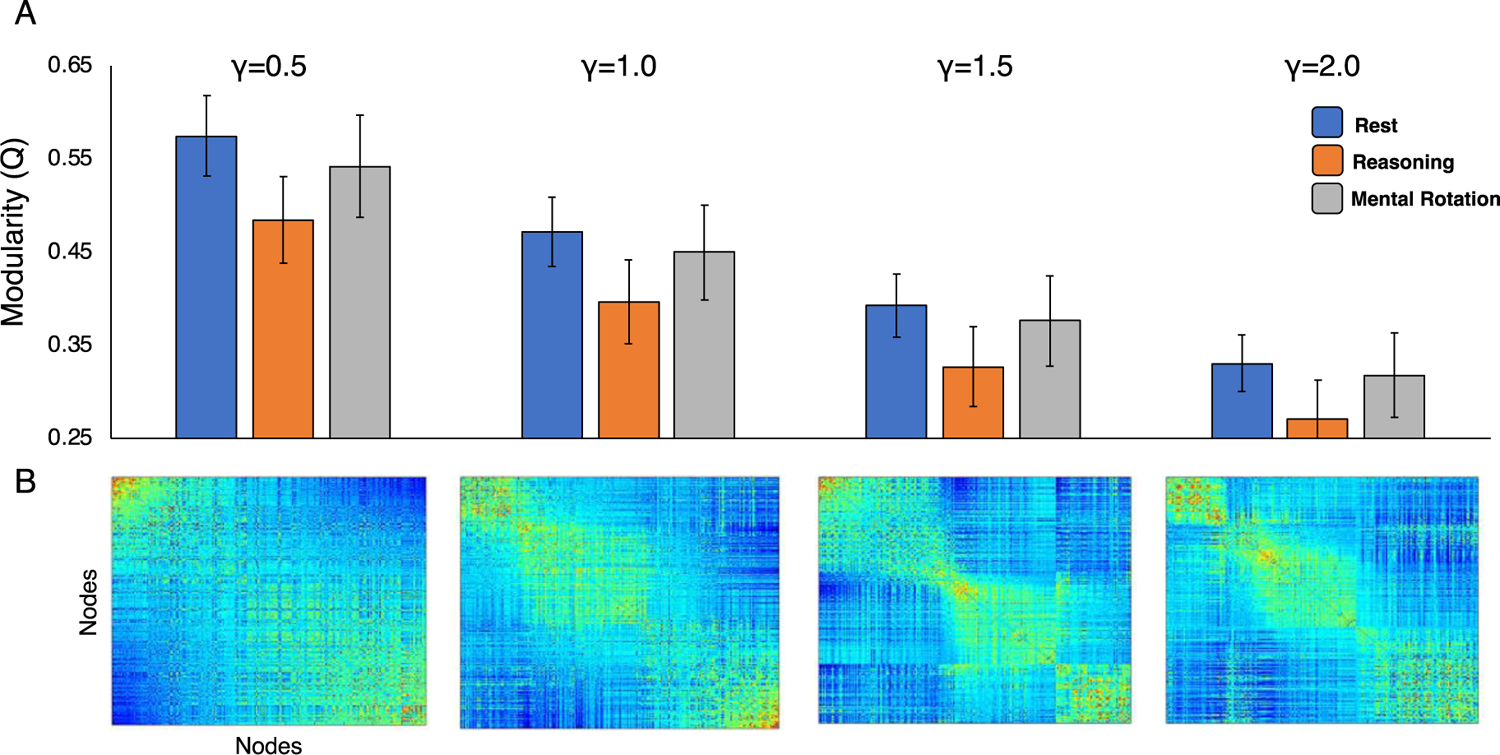
Modularity across resolution parameter and task conditions. Modularity scores (Q) were obtained over a range of resolution parameters for rest, reasoning, and mental rotation. The inverse relationship between Q and *γ* is demonstrated. (A) Q highest during rest relative to both task conditions, with the largest differences observed at lower resolution parameters (γ). (B) Average Q for full sample during rest; increasing *γ* yields more, smaller communities. Community assignments for each node in SI.

Because of the observed differences across γ, we next explored whether task-based reconfigurations in modularity varied based on the resolution parameter. In other words, is resting-state modularity more similar to task-based modularity at different γ? A two factor repeated measures ANOVA indicated that this was the case; the model revealed a significant 4 (Resolution Parameter: 0.5, 1.0, 1.5, 2.0) X 2 (Scan Condition: Rest vs Task) interaction for both reasoning (*F* = 43.50, *p* < 0.0001) and MRT (*F* = 17.62, *p* < 0.0001). Thus, modularity was lower during task conditions than rest, but this effect was the largest when the modularity maximization function was biased towards fewer and larger communities (i.e., at lower γ).

We next investigated whether Q_Rest_ was correlated with Q_Reasoning_ or Q_MRT_ to assess correspondence between resting-state and task-based modularity (Table 1). Across all four resolution parameters, resting-state modularity was unassociated with Q_Reasoning_ (all *r* < 0.12, *p* > 0.34) or Q_MRT_ (all *r* < 0.15, *p* > 0.23). By contrast, modularity during the two task conditions was strongly correlated at all levels of γ (all *r* > 0.34, *p* < 0.005). A post-hoc seemingly unrelated estimate (suest) tested indicated that the association between Q_Reasoning_ and Q_MRT_ did not vary across γ (Q_γ=1.5_ vs. Q_γ=2.0_: χ^2^ = 3.65, *p* = 0.06; all other comparisons: χ^2^ < 1.60, *p* > 0.20).

**Table 1.** Correlations between modularity, GPA, and PSAT scores

| | $Q_{Rest}$<br>$\gamma = 0.5$ | $Q_{Rest}$<br>$\gamma = 1.0$ | $Q_{Rest}$<br>$\gamma = 1.5$ | $Q_{Rest}$<br>$\gamma = 2.0$ | $Q_{Reasoning}$<br>$\gamma = 0.5$ | $Q_{Reasoning}$<br>$\gamma = 1.0$ | $Q_{Reasoning}$<br>$\gamma = 1.5$ | $Q_{Reasoning}$<br>$\gamma = 2.0$ | $Q_{MRT}$<br>$\gamma = 0.5$ | $Q_{MRT}$<br>$\gamma = 1.0$ | $Q_{MRT}$<br>$\gamma = 1.5$ | $Q_{MRT}$<br>$\gamma = 2.0$ | GPA |
| --- | --- | --- | --- | --- | --- | --- | --- | --- | --- | --- | --- | --- | --- |
| $Q_{Rest} \gamma = 1.0$ | 0.91** | -- | | | | | | | | | | | |
| $Q_{Rest} \gamma = 1.5$ | 0.78** | 0.94** | -- | | | | | | | | | | |
| $Q_{Rest} \gamma = 2.0$ | 0.73** | 0.88** | 0.95** | -- | | | | | | | | | |
| $Q_{Reasoning} \gamma = 0.5$ | 0.12 | 0.05 | 0.05 | 0.04 | -- | | | | | | | | |
| $Q_{Reasoning} \gamma = 1.0$ | 0.12 | 0.08 | 0.08 | 0.08 | 0.96** | -- | | | | | | | |
| $Q_{Reasoning} \gamma = 1.5$ | 0.12 | 0.09 | 0.10 | 0.11 | 0.90** | 0.97** | -- | | | | | | |
| $Q_{Reasoning} \gamma = 2.0$ | 0.07 | 0.06 | 0.08 | 0.10 | 0.85** | 0.92** | 0.96** | -- | | | | | |
| $Q_{MRT} \gamma = 0.5$ | 0.07 | 0.05 | 0.05 | 0.03 | 0.37* | 0.37* | 0.35* | 0.41** | -- | | | | |
| $Q_{MRT} \gamma = 1.0$ | 0.14 | 0.11 | 0.11 | 0.10 | 0.37* | 0.38* | 0.37* | 0.41** | 0.95** | -- | | | |
| $Q_{MRT} \gamma = 1.5$ | 0.12 | 0.10 | 0.12 | 0.12 | 0.39* | 0.39* | 0.39* | 0.43** | 0.91** | 0.98** | -- | | |
| $Q_{MRT} \gamma = 2.0$ | 0.10 | 0.10 | 0.13 | 0.13 | 0.42** | 0.43** | 0.42** | 0.47** | 0.89** | 0.95** | 0.98** | -- | |
| GPA | 0.04 | 0.00 | 0.03 | -0.05 | 0.45** | 0.52** | 0.47** | 0.44** | 0.26‡ | 0.23 | 0.24‡ | 0.26‡ | -- |
| PSAT | -0.10 | -0.10 | -0.04 | -0.09 | 0.28‡ | 0.30‡ | 0.27‡ | 0.25 | 0.23 | 0.13 | 0.13 | 0.14 | 0.60** |

**Note:** \*\* $p < 0.001$ , \* $p < 0.01$ , ‡ $p < 0.05$ , uncorrected

Taken together, these results indicate that task demands during reasoning and MRT led to a decrease in modular organization – with the greatest effects for γ ≤ 1.0. Modularity during the completion of both tasks showed little correspondence to modularity at rest, but the strong correlations between Q_Reasoning_ and Q_MRT_ suggest that individuals may exhibit similar modular organization across different task conditions.

### Relationship between modularity and task performance

We next investigated whether individual differences in modularity were related to variation in task performance. Pairwise correlations indicated that – across all levels of *γ* – Q_Reasoning_ was unassociated with reasoning task performance (all *r* < 0.17, *p* > 0.18; Table S3). For MRT, however, we observed a weak, positive association between modularity and MRT performance across all levels of *γ* (0.20 < *r* < 0.28; 0.02 < *p* < 0.11; Table S3).

Zero-order correlations between brain and behavior are fairly common for fMRI studies of individual differences (Power et al., 2012, 2015). However, to obtain a more precise estimate of the relationship between modularity and task performance, we next ran a linear multiple regression model to predict MRT performance in which we controlled for a number of additional variables. First, the model included covariates for socioeconomic status (SES; estimated via mother’s education), race, and gender because functional connectivity has been previously shown to vary by these demographic factors (Satterthwaite et al., 2015; Sripada et al., 2014; Tomasi & Volkow, 2012; Ursache & Noble, 2016; Zhang et al., 2018). We also controlled for participant-level mean framewise displacement to account for in-scanner head movement, which can have substantial impacts on estimates of functional connectivity (Ciric et al., 2017, 2018; Cole et al., 2014; Satterthwaite et al., 2013; Satterthwaite et al., 2012) and differentially affect different brain networks (Van Dijk et al., 2012), even after common denoising pipelines (Ciric et al., 2017, 2018; Van Dijk et al., 2012). There is also evidence to suggest that in-scanner movement may be associated with a wide range of behavioral and demographic measures, including fluid intelligence and verbal ability (Siegel et al., 2017). Finally, we sought to control for lower-level network features that were used to calculate modularity but were not of central interest to the hypothesized relationship between modularity and behavior. To this end, we included the average functional connectivity strength of negative edges as an additional covariate regressor. Note that because positive and negative connections were strongly, inversely related to each other, it would have been inappropriate to include both in the linear regression models. Correlations between modularity and several lower level network features are indicated in Table S2.

In this regression model (for modularity at γ = 1, see Table S4 for similar findings at other resolution parameters), Q_MRT_ was no longer a significant predictor of MRT performance (*β* = −0.12, *p* = 0.56), but mean framewise displacement was (*β* = −0.41, *p* = 0.003). Thus, participants who moved less in the scanner performed better on the task. These results suggest that the bivariate association between Q_MRT_ and MRT performance may actually reflect differences in framewise displacement, consistent with recent work reporting associations between movement and cognitive performance (Siegel et al., 2017). That is, because framewise displacement was negatively correlated with both Q (*r* = −0.30, *p* = 0.009) and MRT performance (i.e., less movement associated with better performance; *r* = −0.28, *p* = 0.02), Q_MRT_ was positively correlated with performance. Broadly, these results highlight the importance of controlling for in-scanner movement to more precisely estimate the relationship between brain and behavior.

### Relationship between modularity and academic learning and ability

Given prior work establishing a positive association between modularity and intelligence (Barbey, 2018; Chiappe & MacDonald, 2005; Finn et al., 2015; Hilger et al., 2017, 2020, see Kruschwitz et al., 2018 for alternative account) and learning (Arnemann et al., 2015; Baniqued et al., 2018; Duncan & Small, 2016; Gallen et al., 2016; Gallen & D’Esposito, 2019), we examined whether modularity was related to PSAT scores (i.e., academic ability) and GPA (i.e., a proxy for academic learning).

We found no evidence of an association between resting-state modularity and academic ability; PSAT scores were uncorrelated with Q_Rest_ at all resolution parameters (all *r* < 0.05, *p* > 0.41; Table 1). Similarly, student GPA – an broad index of academic learning (Duff et al., 2004; Komarraju et al., 2011) – was uncorrelated with Q_Rest_ (all *r* < 0.04, *p* > 0.69; Table 1, Fig. 3). These findings run counter to perspectives that suggest that modularity – especially modularity during resting-state fMRI – may be a biomarker of learning or intelligence (Barbey, 2018; Gallen & D’Esposito, 2019), but do adhere with a large sample study (N = 1096) in which the authors found no association between global network measures of resting-state functional connectivity and intelligence (Kruschwitz et al., 2018).

**Figure 3.**
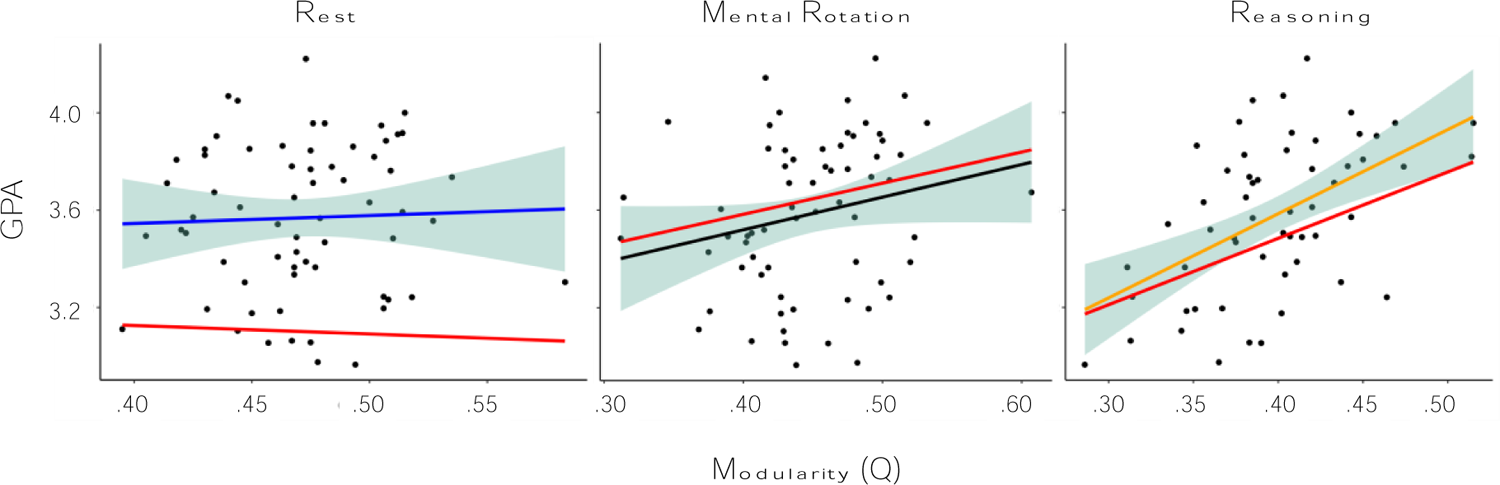
Associations between modularity and GPA. All plots display results at γ = 1.0. For all plots, red line indicates standardize beta for Q in multiple regression model. No association between GPA and Q during rest, but a positive correlation was observed for both task conditions. Note that, while QMRT was not significantly associated with GPA (*p* = 0.33), the magnitude of the effect was similar as observed in the zero-order correlations. 95% CI indicated for zero-order correlations. See Tables S6, S7 for results across all resolution parameters.

Although resting-state modularity was unassociated with PSAT scores or GPA, we next investigated whether academic ability or learning was associated with *task*-*based* modularity. Pairwise correlations revealed a positive correlation between Q_Reasoning_ and academic ability (PSAT scores; γ = 0.5 to 1.5: all *r* > 0.26, *p* < 0.05; γ = 2.0: *r* = 0.25, *p* = 0.06), as well as an even stronger association between Q_Reasoning_ and GPA (i.e., a proxy for academic learning; all *r* > 0.43, *p* < 0.0007; Table 1, Fig. 3). We also observed a nominally positive – although non-significant – association between Q_MRT_ and PSAT scores (all *r* > 0.12, *p* > 0.07, Table 1). Q_MRT_ was also correlated with GPA across all resolution parameters (all *r* > 0.23, *p* < 0.06; Table 1, Fig. 3).

Given evidence that task-based modularity was associated with estimates of academic ability and learning but resting-state modularity was not, we next asked whether this difference was significant. That is, were Q_Reasoning_ or Q_MRT_ significantly more correlated with PSAT scores and GPA than was Q_Rest_? Another series of suest tests indicated that Q_Reasoning_ was significantly more correlated with both GPA (χ^2^ = 15.78, *p* = 0.0001) and PSAT scores (χ^2^ = 8.35, *p* = 0.004) than was Q_Rest_ (results reflect Q at γ=1; results at other resolution parameters indicated in Table S5). Q_MRT_ was also more strongly correlated with GPA and PSAT scores than was Q_Rest_ at a trend level (GPA: χ^2^ = 2.91, *p* = 0.09; PSAT: χ^2^ = 2.67, *p* = 0.10).

Lastly, we ran a series of additional linear multiple regression models using the same covariates as those previously described to predict GPA and PSAT scores (i.e., demographic information, task-specific framewise displacement and functional connectivity strength of negative edges). Note that regression models reported here examined modularity for γ = 1 (see Table S6, S7 for results at other resolution parameters).

Regression models indicated that Q_MRT_ was no longer associated with PSAT scores (*β* = − 0.09, *p* = 0.72) or GPA (*β* = 0.23, *p* = 0.33; Table S7). Post-hoc analyses, however, indicated that the effect size of Q_MRT_ in the regression model to predict GPA was not significantly different than that observed in the zero-order correlations (χ^2^ < 0.01, *p* = 0.97). Thus, the loss of significance likely stems from fewer degrees of freedom in the multiple regression model. Q_Reasoning_ was not a significant predictor of PSAT scores in a multiple regression model (*β* = −0.16, *p* = 0.46), but remained a significant predictor of GPA (*β* = 0.42, *p* = 0.04; Table S6, Fig. 3). Taken together, these results indicate that modularity – particularly during task conditions – may be associated with GPA, a broad index of academic learning.

## Discussion

In this study, we examined modularity across different resolution parameters and task conditions, as well as the extent to which modularity related to task performance and indices of academic learning and ability. We found that modularity was higher at rest than during task conditions, and that the magnitude of this difference was largest when modularity calculations were biased towards larger and fewer communities (i.e., *γ* ≤ 1.0). Results further indicated that Q_Rest_ was unassociated with task-based modularity. That is, there was little correspondence between resting-state modularity and modular organization when completing cognitive tasks. By contrast, we observed strong correlations between Q_MRT_ and Q_Reasoning_, suggesting that individuals may exhibit similar levels of modularity across different task conditions. Individual differences in modularity were unrelated to task performance, but showed associations with GPA as a proxy for academic learning (participants with higher modularity during both task conditions had higher GPAs).

Modular organization, the clustering of network nodes into multiple and distinct subgraphs (Genon et al., 2018; Lorenz et al., 2017; Newman, 2006; Power et al., 2011; Sporns & Betzel, 2016; Yeo et al., 2011; Toga et al., 2006), is thought to allow for flexible adaptation and integration through the rearrangement of pertinent subnetworks in response to task demands (Kashtan et al., 2007; Kashtan & Alon, 2005; Wig, 2017). That is, tasks induce a reconfiguration of intrinsic functional modules. If this is true, then one would expect to see a reduction in brain-wide modularity when an individual is engaged in a task because otherwise-segregated subgraphs must communicate in order to execute the relevant goals or objectives. Indeed, task-based reductions in modularity have been frequently reported (e.g., Cohen & D’Esposito, 2016; Di et al., 2013; Finc et al., 2017, 2020; Kitzbichler et al., 2011). The present study provides further support for task-induced decreases in modularity; modularity was higher at rest than during reasoning or MRT.

Brains are hierarchical (Bassett & Bullmore, 2006; Betzel et al., 2017; Betzel & Bassett, 2016; Reichardt & Bornholdt, 2006), meaning that the community organization at one level is nested within larger levels of organization. For example, at a coarse level, one could consider the two hemispheres to reflect discrete communities. Moving down to finer granularity, modules are reflected in separable functional networks (e.g., frontoparietal network), smaller sub-networks (frontal, parietal), gray matter nuclei, and even cell columns (Sporns, 2010). Therefore, it is important to estimate global network statistics (i.e., modularity) over different scales because organizational properties may be meaningfully different for larger communities (i.e., at lower values of the resolution parameter, *γ*) than for smaller communities contained within them (i.e., at higher values of *γ*). Intriguingly, the extent of the above-mentioned task-based decreases in modularity varied as a function of community size; we observed larger differences between rest and the two task conditions when estimating the modular organization of larger communities compared to the differences observed for smaller communities. The modularity statistic (Q) is positively associated with community size (i.e., Q is larger when derived from larger communities), thus the discrepancy between rest and task conditions shrunk alongside decreasing modularity. That is, differences between Q_Rest_ and Q_Task_ were less pronounced for smaller communities. This result may stem from the finding that resting-state subgraph intercommunication is already high when estimating the network organization of smaller communities (as evidenced by lower Q). Task demands are associated decreased segregation of resting-state functional networks (Cole et al., 2014; Medaglia et al., 2015; Power et al., 2011; Yeo et al., 2011), but the relative extent of this departure is less when the intrinsic communities are already interconnected. Said differently, results of the present study indicate that as one moves further down the hierarchy of functional organization in the brain, task-induced changes in modularity decrease in magnitude.

This is not to say, however, that alterations (or lack thereof) to smaller communities are meaningless. Rather, in order to quantify how a given task influences the organization of functional brain networks, it is necessary to identify the resolution at which such effects should be observed. By moving up and down the brain’s organizational hierarchy, we become better able to make such determinations. The present findings suggest that that brain’s modular organization at finer scales is fairly robust to external demands induced by cognitive tasks, whereas coarser communities are more subject to task-based reorganization. Future network neuroscience work should keep such considerations in mind when examining associations between community organization and other variables of interest.

Consistent with the putative relationship between modularity and adaptability (Anderson & Finlay, 2014; Bassett & Bullmore, 2006; Mattar et al., 2018; Meunier et al., 2009), a growing number of studies have indicated that modularity may be a biomarker of learning (Arnemann et al., 2015; Baniqued et al., 2018, 2019; Chaddock-Heyman et al., 2020; Duncan & Small, 2016; Gallen et al., 2016; Gallen & D’Esposito, 2019). Results from the present study lend some support to these perspectives. Most notably, we observed a significant positive association between task-based modularity and GPA – a reasonable proxy for academic learning (Duff et al., 2004; Komarraju et al., 2011). Even after controlling for a number of potentially confounding variables, including demographic information, in-scanner movement, and other functional network characteristics, the association between modularity during the reasoning task and GPA remained significant. For MRT, although modularity was no longer significantly predictive of GPA, the magnitude of the effect size in the multiple regression model was not significantly different than in the zero-order correlations, suggesting that the reduction in significance stemmed from fewer degrees of freedom.

The present findings differ from past work in a number of ways. First, to our knowledge, the extended classroom-based learning in the present study is qualitatively distinct from prior work examining modularity and learning, which has been restricted to comparatively shorter interventions. GPA – used in the present study as a proxy for academic learning – is subject to a host of additional factors including motivation, attendance, and effort (Goldman & Widawski, 1976; Noble & Sawyer, 2004; Stiggins et al., 1989). On the one hand, this complicates the specificity of the present findings as the observed effects could be attributed to any combination of these variables (and several others). However, given the influence of high school GPA on later life achievement (French et al., 2015; Noble & Sawyer, 2004; Noble & Sawyer, 2002; Sawyer, 2013; Schulenberg et al., 1994), identifying its neural correlates is likely to be of interest to educators and policy-makers, and may have additional value if it can provide insights beyond what can be obtained through behavioral assessments alone (Bruer, 1997; Goswami, 2009; Supekar et al., 2013). More broadly, to better understand the relationship between brain organization and learning, this association should be investigated for contexts in which learning matters most (i.e., in the real-world and schools).

Another novel contribution of the present study is that the strongest associations between learning and modularity were obtained during reasoning and MRT; GPA was strongly associated with task-based modularity but showed no relationship with modularity during the resting scan. These findings run counter to a body of work that has identified associations between resting-state modularity and learning (Arnemann et al., 2015; Baniqued et al., 2018, 2019; Chaddock-Heyman et al., 2020; Duncan & Small, 2016; Gallen et al., 2016; Gallen & D’Esposito, 2019). Because modularity was higher at rest than during either of the task conditions, our findings suggest that additional insights may be gained by considering the extent to which one is able to maintain intrinsic, resting-state modularity *despite* competing task demands. Note, however, that additional analyses involving a “difference score” (i.e., task-based modularity subtracted from resting-state modularity; see SI) were redundant with those obtained using task-based modularity alone. Therefore, it is unclear whether the amount of change from intrinsic (i.e., resting-state) functional organization can offer unique insights beyond modular organization during the task itself. Regardless, the present findings indicate that task-based modularity may be a better candidate neural biomarker of academic learning than modularity at rest, although it remains important to control for demographics, in-scanner motion, and lower-level network features.

We also examined associations between modularity and PSAT scores. As noted above, GPA is likely to reflect a number of factors including academic ability and intelligence, but it also clearly relates to learning as well (Duff et al., 2004; Komarraju et al., 2011). Learning is less related to PSAT scores, which are generally believed to reflect academic ability and fluid intelligence (Coyle, 2015; Lent et al., 1986). In contrast to results involving GPA, modularity was only weakly associated with PSAT scores. This result challenges prior research that has associated modular organization with assessments of general intelligence (i.e., the Wechsler Abbreviated Scale of Intelligence; Hilger et al., 2017, 2020; Weschler, 2011) and memory (Stanley et al., 2014; Stevens et al., 2012; Wig, 2017). At least one recent study, however, concluded that resting-state functional connectivity shows little correspondence with intelligence (Kruschwitz et al., 2018). It is plausible – although speculative – that these discrepant findings could be due to publication bias: research groups that identify associations between modularity and intelligence publish their results, but researchers who fail to identify such relationships in their own datasets do not. Indeed, many leading scientific publishers have specifically highlighted the value of publishing – rather than effectively burying – “null” or “negative” results (“Rewarding Negative Results Keeps Science on Track,” 2017). Given the increased use of graph theoretic techniques to examine the brain bases of multifaceted psychological constructs (e.g., Barbey, 2018; Bassett & Mattar, 2017; Gallen & D’Esposito, 2019), there is an especially pressing need for transparent reporting within network neuroscience. Mitigating publication bias has been shown to improve replicability (Dwan et al., 2008; Szucs & Ioannidis, 2017), especially with smaller samples (Button et al., 2013).

One notable limitation of the present study is that the resting scan was only five minutes in duration. Although this is reasonably consistent with prior work that has related modularity to intervention-related gains (e.g., Arnemann et al., 2015; Baniqued et al., 2018; Gallen et al., 2016), the effect of scan duration can impact functional connectivity estimates. Specifically, scans of 12 minutes or longer are substantially more reliable than 5-7 minute scans (Birn et al., 2013). Because both task conditions *were* of a sufficient duration to allow for stabilization of functional networks (Van Dijk et al., 2009), it is plausible that results from a longer resting-state scan would have yielded comparable results to the task conditions.

To conclude, the present study investigated modular organization at rest and during two task conditions. We observed higher modularity at rest than during task, with the most pronounced differences for larger functional communities. We also investigated the extent to which modularity related to real-world learning (i.e., in schools), extending prior work that has relied on shorter and/or lab-based training paradigms. Although resting-state modularity was unassociated with academic learning, we found that participants with higher task-based modularity had higher GPAs. Future work should continue to explore the implications of brain network organization on learning in real-world context, and should pay particular attention to differential associations of resting vs. task-related connectivity.

## Acknowledgements

The authors thank Charles Lynch for his help conducting and conceptualizing analyses. They also thank Ian Lyons and Chandan Vaidya for their insightful comments during the writing of this manuscript.

## Funding

This research was funded by a grant from the John Templeton Foundation to A.B.W. and A.E.G. [ID 61114], and grants to A.E.G. from the National Science Foundation [DRL-1420481, EHR-1661065, EHR-1920682].

## Author Contributions

**A.B.W.:** Conceptualization, Methodology, Formal analysis, Investigation, Writing – Original draft, Writing – Review & editing, Visualization, Funding acquisition. **R.A.C.:** Formal analysis, Writing – Review & editing. **R.F.B.:** Writing – Review & editing, Supervision. **A.E.G.:** Writing – Review & editing, Supervision, Project administration, Funding acquisition.

## Notes

https://osf.io/naj3y/

### Competing Interest Statement

The authors have declared no competing interest.

