## Supplementary Information for "Exploring Functional Brain Network Modularity in Educational Contexts"

**1. Accuracy-RT Composite Score**

Conventionally, speed (RT) and accuracy have both be used to measure performance on cognitive tasks, including tasks of reasoning and mental rotation. While typically consider separately, composite measures integrating RT and accuracy are gaining traction (Lyons et al., 2014; Vandierendonck, 2017; Woltz & Was, 2006). According to a recent review (Vandierendonck, 2017), composite scores of speed and accuracy are productive when both are dependent upon the same cognitive processes, as evident by a significant and positive association between RT and error rate. That is, faster performance should be correlated with greater accuracy. Furthermore, integrated measures are most appropriate when accuracy scores are generally high. While there is still no consensus as to which, if any, variables can best capture variation in RT and accuracy, Rate Correct Score (RCS; Woltz & Was, 2006) has received generally favorable endorsements (Hughes et al., 2014; Vandierendonck, 2017). RCS is calculated as:

$$RCS = \frac{number\ of\ correct\ responses\ of\ considered\ trials}{\sum RT\ of\ considered\ trials}$$

and can be interpreted as number of correct responses per unit of time.

RCS was considered for use in the present study if participants displayed high overall accuracy and a strong, positive correlation between RT and error rate. Results indicated that this was indeed the case. Accuracy scores were high for reasoning (M = 81.4%, SD = 12.4%) and MRT (M = 75.9%, SD = 12.6%). Further, error rate and RT were significantly positively correlated for reasoning ($r = 0.56$, $p < 0.0001$) and MRT ($r = 0.29$, $p = 0.01$). Thus, as described in the main text, RCS was used as the primary variable for performance on both tasks.

**2. Modularity "Difference Score"**

Because modularity scores were higher at rest, we explored whether the extent to which an individual is able to maintain intrinsic modularity (i.e. resting-state modularity) while completing a complex reasoning tasks (reasoning, MRT) was related to academic learning and ability. To do this, we subtracted resting-state modularity from modularity for each of the two task conditions, producing separate modularity difference scores for the two tasks. For Reasoning, the difference score was significantly correlated with GPA ($\gamma$=1.0: $r = 0.46$, $p = 0.0003$; Tables S8) and weakly associated PSAT scores ($\gamma$=1.0: $r = 0.32$, $p = 0.02$; Tables S8). For MRT, the difference score was weakly associated with GPA ($\gamma$=1.0: $r = 0.27$, $p = 0.03$; Tables S8) but not with PSAT scores ($\gamma$=1.0: $r = 0.19$, $p = 0.14$; Tables S8). These values are similar to those obtained from $Q_{Reasoning}$ alone, suggesting subtracting $Q_{Rest}$ does not meaningfully change results reported in the main text.

Reasoning

The Ape is better than the Cat.
The Dog is worse than the Cat.
**The Ape is better than the Dog.**

Mental Rotation

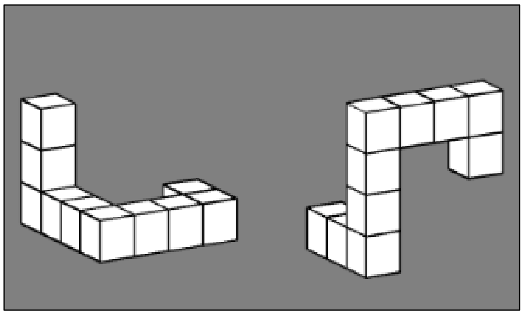

**Figure S1. Sample Reasoning and Mental Rotation Tasks.**

**Table S1. Available data and descriptive statistics for all fMRI conditions**

| | Rest | Reasoning | MRT |
|---|---|---|---|
| **N** | 75 | 66 | 74 |
| **Age** | 16.71 ± 0.49 | 16.73 ± 0.48 | 16.72 ± 0.48 |
| **Gender** | 50.67% Male<br>49.33% Female | 42.42% Male<br>57.58% Female | 48.65% Male<br>51.35% Female |
| **PSAT Available** | 66 | 57 | 65 |
| **GPA Available** | 68 | 59 | 67 |

**Table S2. Associations between modularity and "lower level" network features.**

| | Rest | | | | Reasoning | | | | MRT | | | |
|---|---|---|---|---|---|---|---|---|---|---|---|---|
| | $Q_{\gamma=0.5}$ | $Q_{\gamma=1.0}$ | $Q_{\gamma=1.5}$ | $Q_{\gamma=2.0}$ | $Q_{\gamma=0.5}$ | $Q_{\gamma=1.0}$ | $Q_{\gamma=1.5}$ | $Q_{\gamma=2.0}$ | $Q_{\gamma=0.5}$ | $Q_{\gamma=1.0}$ | $Q_{\gamma=1.5}$ | $Q_{\gamma=2.0}$ |
| Average FC | 0.13 | -0.03 | -0.16 | -0.11 | -0.01 | -0.07 | -0.10 | -0.07 | 0.25 | 0.26 | 0.26 | 0.28 |
| | 0.26 | 0.77 | 0.18 | 0.37 | 0.93 | 0.59 | 0.43 | 0.58 | 0.03 | 0.03 | 0.03 | 0.02 |
| Number Pos. Edges | -0.11 | -0.36 | -0.48 | -0.48 | -0.52 | -0.60 | -0.65 | -0.65 | -0.32 | -0.45 | -0.50 | -0.53 |
| | 0.36 | <.001 | <.001 | <.001 | <.001 | <.001 | <.001 | <.001 | <.001 | <.001 | <.001 | <.001 |
| Average FC Pos. Edges | 0.74 | 0.73 | 0.67 | 0.68 | 0.77 | 0.78 | 0.78 | 0.78 | 0.79 | 0.80 | 0.81 | 0.82 |
| | <.001 | <.001 | <.001 | <.001 | <.001 | <.001 | <.001 | <.001 | <.001 | <.001 | <.001 | <.001 |
| Number Neg. Edges | 0.11 | 0.36 | 0.48 | 0.48 | 0.52 | 0.60 | 0.65 | 0.65 | 0.32 | 0.45 | 0.50 | 0.53 |
| | 0.36 | <.001 | <.001 | <.001 | <.001 | <.001 | <.001 | <.001 | <.001 | <.001 | <.001 | <.001 |
| Average FC Neg. Edges | -0.77 | -0.72 | -0.65 | -0.64 | -0.76 | -0.75 | -0.74 | -0.72 | -0.77 | -0.73 | -0.72 | -0.72 |
| | <.001 | <.001 | <.001 | <.001 | <.001 | <.001 | <.001 | <.001 | <.001 | <.001 | <.001 | <.001 |

**Note**: *r* value indicated on top row, *p* value indicated on bottom row. Average FC of negative edges used as covariate regressor in linear regression models

**Table S3. Zero-order correlations between modularity and task performance**

| | Reasoning (RCS) | MRT (RCS) |
|---|---|---|
| $Q_{\gamma=0.5}$ | 0.14 | 0.28 |
| | 0.279 | 0.023 |
| $Q_{\gamma=1.0}$ | 0.17 | 0.21 |
| | 0.180 | 0.083 |
| $Q_{\gamma=1.5}$ | 0.13 | 0.20 |
| | 0.299 | 0.102 |
| $Q_{\gamma=2.0}$ | 0.15 | 0.25 |
| | 0.239 | 0.046 |

**Note**: *r* value indicated on top row, *p* value indicated on bottom row, uncorrected

**Table S4. Linear regression models to predict MRT performance**

| MRT Performance | $\beta$ | Std. Err. | t | P | 95% CI | |
|---|---|---|---|---|---|---|
| MRT $Q_{\gamma=0.5}$ | 0.02 | 0.20 | 0.10 | 0.919 | -0.39 | 0.43 |
| Mother total education | 0.09 | 0.01 | 0.73 | 0.467 | -0.01 | 0.02 |
| Gender | 0.33 | 0.01 | 2.74 | 0.008 | 0.01 | 0.06 |
| Race/Ethnicity | -0.07 | 0.01 | -0.62 | 0.536 | -0.03 | 0.02 |
| Mean FD | -0.36 | 0.08 | -2.59 | 0.012 | -0.35 | -0.05 |
| Strength Negative FC | -0.20 | 0.83 | -0.92 | 0.361 | -2.41 | 0.89 |
| MRT $Q_{\gamma=1.0}$ | -0.12 | 0.20 | -0.59 | 0.558 | -0.52 | 0.28 |
| Mother total education | 0.07 | 0.01 | 0.58 | 0.564 | -0.01 | 0.01 |
| Gender | 0.32 | 0.01 | 2.66 | 0.01 | 0.01 | 0.06 |
| Race/Ethnicity | -0.08 | 0.01 | -0.73 | 0.469 | -0.04 | 0.02 |
| Mean FD | -0.41 | 0.07 | -3.07 | 0.003 | -0.37 | -0.08 |
| Strength Negative FC | -0.32 | 0.75 | -1.60 | 0.116 | -2.70 | 0.30 |
| MRT $Q_{\gamma=1.5}$ | -0.17 | 0.20 | -0.87 | 0.388 | -0.57 | 0.23 |
| Mother total education | 0.07 | 0.01 | 0.56 | 0.58 | -0.01 | 0.01 |
| Gender | 0.32 | 0.01 | 2.69 | 0.009 | 0.01 | 0.06 |
| Race/Ethnicity | -0.09 | 0.01 | -0.77 | 0.443 | -0.04 | 0.02 |
| Mean FD | -0.43 | 0.07 | -3.24 | 0.002 | -0.38 | -0.09 |
| Strength Negative FC | -0.36 | 0.71 | -1.87 | 0.067 | -2.75 | 0.09 |
| MRT $Q_{\gamma=2.0}$ | -0.04 | 0.21 | -0.23 | 0.821 | -0.48 | 0.38 |
| Mother total education | 0.09 | 0.01 | 0.68 | 0.497 | -0.01 | 0.02 |
| Gender | 0.33 | 0.01 | 2.72 | 0.009 | 0.01 | 0.06 |
| Race/Ethnicity | -0.08 | 0.01 | -0.67 | 0.504 | -0.04 | 0.02 |
| Mean FD | -0.39 | 0.07 | -2.92 | 0.005 | -0.35 | -0.07 |
| Strength Negative FC | -0.26 | 0.71 | -1.36 | 0.181 | -2.38 | 0.46 |

**Note**: Results for $\gamma$ = 1.0 reported in main text; mean FD = mean framewise displacement

**Table S4. Comparison of effect sizes of reasoning and MRT (to rest) for GPA and PSAT Scores**

| | **Reasoning** | | **MRT** | |
|---|---|---|---|---|
| | GPA | PSAT Scores | GPA | PSAT Scores |
| $Q_{\gamma=0.5}$ | Reasoning > Rest<br>$\chi^2 = 8.07$; $p = 0.005$ | Reasoning > Rest<br>$\chi^2 = 6.96$; $p = 0.008$ | MRT > Rest<br>$\chi^2 = 2.08$; $p = 0.149$ | MRT > Rest<br>$\chi^2 = 4.14$; $p = 0.042$ |
| $Q_{\gamma=1.0}$ | Reasoning > Rest<br>$\chi^2 = 15.78$, p = 0.0001 | Reasoning > Rest<br>$\chi^2 = 8.35$, p = 0.004 | MRT > Rest<br>$\chi^2 = 2.921$, p = 0.088 | MRT > Rest<br>$\chi^2 = 2.67$, p = 0.102 |
| $Q_{\gamma=1.5}$ | Reasoning > Rest<br>$\chi^2 = 10.07$; $p = 0.002$ | Reasoning > Rest<br>$\chi^2 = 4.86$; $p = 0.028$ | MRT > Rest<br>$\chi^2 = 2.75$; $p = 0.097$ | MRT > Rest<br>$\chi^2 = 1.74$; $p = 0.187$ |
| $Q_{\gamma=2.0}$ | Reasoning > Rest<br>$\chi^2 = 11.18$; $p = 0.001$ | Reasoning > Rest<br>$\chi^2 = 5.29$; $p = 0.022$ | MRT > Rest<br>$\chi^2 = 6.15$; $p = 0.013$ | MRT > Rest<br>$\chi^2 = 2.78$; $p = 0.095$ |

**Note**: Results for $\gamma = 1.0$ reported in main text. Values calculated from seemingly unrelated estimate test (suest)

**Table S5. Linear regression models (for all γ) to predict GPA and PSAT from $Q_{Reasoning}$**

| | GPA Regression Models | | | | | PSAT Regression Models | | | | |
|---|---|---|---|---|---|---|---|---|---|---|
| | **B** | **Std. Err.** | **t** | **P** | ***β*** | **B** | **Std. Err.** | **t** | **P** | ***β*** |
| Reasoning $Q_{\gamma=0.5}$ | 1.58 | 1.34 | 1.18 | 0.244 | 0.24 | -97.33 | 99.85 | -0.97 | 0.335 | -0.21 |
| Mother total education | 0.05 | 0.04 | 1.22 | 0.226 | 0.16 | 5.79 | 3.02 | 1.92 | 0.061 | 0.27 |
| Gender | -0.05 | 0.08 | -0.59 | 0.558 | -0.08 | -9.18 | 6.33 | -1.45 | 0.154 | -0.20 |
| Race/Ethnicity | 0.05 | 0.09 | 0.52 | 0.607 | 0.06 | -2.24 | 6.62 | -0.34 | 0.737 | -0.04 |
| Mean FD | -0.84 | 0.47 | -1.78 | 0.082 | -0.25 | -49.51 | 35.03 | -1.41 | 0.164 | -0.21 |
| Strength Negative FC | -3.60 | 7.86 | -0.46 | 0.649 | -0.09 | -1298.54 | 585.79 | -2.22 | 0.031 | -0.45 |
| Reasoning $Q_{\gamma=1.0}$ | 2.81 | 1.35 | 2.08 | 0.043 | 0.42 | -76.63 | 103.31 | -0.74 | 0.462 | -0.16 |
| Mother total education | 0.05 | 0.04 | 1.17 | 0.246 | 0.15 | 5.74 | 3.04 | 1.89 | 0.065 | 0.27 |
| Gender | -0.03 | 0.08 | -0.42 | 0.679 | -0.05 | -8.90 | 6.37 | -1.40 | 0.169 | -0.20 |
| Race/Ethnicity | 0.05 | 0.09 | 0.57 | 0.569 | 0.07 | -2.32 | 6.66 | -0.35 | 0.729 | -0.04 |
| Mean FD | -0.60 | 0.48 | -1.24 | 0.223 | -0.18 | -50.72 | 36.89 | -1.37 | 0.176 | -0.21 |
| Strength Negative FC | 0.46 | 7.41 | 0.06 | 0.95 | 0.01 | -1178.28 | 566.16 | -2.08 | 0.043 | -0.41 |
| Reasoning $Q_{\gamma=1.5}$ | 1.95 | 1.41 | 1.38 | 0.173 | 0.28 | -104.36 | 105.31 | -0.99 | 0.327 | -0.21 |
| Mother total education | 0.05 | 0.04 | 1.30 | 0.199 | 0.17 | 5.65 | 3.01 | 1.88 | 0.067 | 0.26 |
| Gender | -0.05 | 0.08 | -0.61 | 0.547 | -0.08 | -9.03 | 6.29 | -1.44 | 0.158 | -0.20 |
| Race/Ethnicity | 0.05 | 0.09 | 0.52 | 0.607 | 0.06 | -2.23 | 6.62 | -0.34 | 0.737 | -0.04 |
| Mean FD | -0.72 | 0.49 | -1.47 | 0.148 | -0.22 | -54.59 | 36.79 | -1.48 | 0.144 | -0.23 |
| Strength Negative FC | -3.37 | 7.38 | -0.46 | 0.65 | -0.08 | -1256.39 | 551.08 | -2.28 | 0.027 | -0.43 |
| Reasoning $Q_{\gamma=2.0}$ | 1.52 | 1.40 | 1.08 | 0.283 | 0.21 | -110.08 | 104.50 | -1.05 | 0.297 | -0.21 |
| Mother total education | 0.05 | 0.04 | 1.30 | 0.201 | 0.17 | 5.70 | 3.01 | 1.89 | 0.064 | 0.26 |
| Gender | -0.06 | 0.08 | -0.67 | 0.504 | -0.09 | -9.08 | 6.28 | -1.45 | 0.155 | -0.20 |
| Race/Ethnicity | 0.05 | 0.09 | 0.52 | 0.604 | 0.07 | -2.37 | 6.62 | -0.36 | 0.721 | -0.05 |
| Mean FD | -0.81 | 0.49 | -1.66 | 0.103 | -0.24 | -53.81 | 36.10 | -1.49 | 0.143 | -0.22 |
| Strength Negative FC | -5.21 | 7.19 | -0.72 | 0.472 | -0.13 | -1250.94 | 532.58 | -2.35 | 0.023 | -0.43 |

Note: Results for $\gamma$ = 1.0 reported in main text

**Table S6. Linear regression models (for all γ) to predict GPA and PSAT from $Q_{MRT}$**

| | GPA Regression Models | | | | | PSAT Regression Models | | | | |
|---|---|---|---|---|---|---|---|---|---|---|
| | **B** | **Std. Err.** | **t** | **P** | ***β*** | **B** | **Std. Err.** | **t** | **P** | ***β*** |
| MRT $Q_{\gamma=0.5}$ | 1.94 | 1.39 | 1.40 | 0.168 | 0.35 | 101.98 | 106.30 | 0.96 | 0.341 | 0.26 |
| Mother total education | 0.04 | 0.04 | 1.08 | 0.283 | 0.15 | 4.47 | 2.89 | 1.55 | 0.127 | 0.22 |
| Gender | 0.01 | 0.08 | 0.11 | 0.915 | 0.01 | -4.38 | 6.28 | -0.70 | 0.489 | -0.10 |
| Race/Ethnicity | 0.04 | 0.09 | 0.49 | 0.628 | 0.06 | 1.33 | 6.65 | 0.20 | 0.842 | 0.03 |
| Mean FD | -0.45 | 0.52 | -0.87 | 0.387 | -0.14 | 17.59 | 38.74 | 0.45 | 0.652 | 0.07 |
| Strength Negative FC | 3.56 | 5.61 | 0.63 | 0.529 | 0.16 | 20.18 | 423.97 | 0.05 | 0.962 | 0.01 |
| MRT $Q_{\gamma=1.0}$ | 1.35 | 1.38 | 0.98 | 0.333 | 0.23 | -38.05 | 105.19 | -0.36 | 0.719 | -0.09 |
| Mother total education | 0.04 | 0.04 | 1.06 | 0.294 | 0.15 | 3.91 | 2.94 | 1.33 | 0.188 | 0.19 |
| Gender | 0.01 | 0.08 | 0.08 | 0.936 | 0.01 | -5.63 | 6.35 | -0.89 | 0.379 | -0.12 |
| Race/Ethnicity | 0.04 | 0.09 | 0.40 | 0.689 | 0.05 | 0.14 | 6.69 | 0.02 | 0.983 | 0.00 |
| Mean FD | -0.61 | 0.50 | -1.23 | 0.225 | -0.18 | -11.24 | 37.15 | -0.30 | 0.763 | -0.05 |
| Strength Negative FC | 0.98 | 5.15 | 0.19 | 0.849 | 0.04 | -448.95 | 386.94 | -1.16 | 0.251 | -0.28 |
| MRT $Q_{\gamma=1.5}$ | 1.42 | 1.38 | 1.03 | 0.306 | 0.23 | -39.55 | 103.06 | -0.38 | 0.703 | -0.09 |
| Mother total education | 0.04 | 0.04 | 1.04 | 0.301 | 0.15 | 3.94 | 2.92 | 1.35 | 0.182 | 0.19 |
| Gender | 0.00 | 0.08 | 0.04 | 0.965 | 0.01 | -5.53 | 6.30 | -0.88 | 0.384 | -0.12 |
| Race/Ethnicity | 0.04 | 0.09 | 0.41 | 0.683 | 0.05 | 0.09 | 6.70 | 0.01 | 0.99 | 0.00 |
| Mean FD | -0.61 | 0.49 | -1.23 | 0.223 | -0.18 | -11.17 | 36.47 | -0.31 | 0.761 | -0.05 |
| Strength Negative FC | 0.91 | 4.90 | 0.19 | 0.853 | 0.04 | -444.74 | 362.67 | -1.23 | 0.225 | -0.27 |
| MRT $Q_{\gamma=2.0}$ | 1.84 | 1.46 | 1.26 | 0.214 | 0.27 | -43.87 | 109.90 | -0.40 | 0.691 | -0.09 |
| Mother total education | 0.04 | 0.04 | 1.05 | 0.299 | 0.15 | 3.98 | 2.90 | 1.37 | 0.176 | 0.20 |
| Gender | 0.00 | 0.08 | 0.05 | 0.96 | 0.01 | -5.52 | 6.30 | -0.88 | 0.384 | -0.12 |
| Race/Ethnicity | 0.04 | 0.09 | 0.44 | 0.665 | 0.06 | 0.09 | 6.69 | 0.01 | 0.989 | 0.00 |
| Mean FD | -0.56 | 0.49 | -1.15 | 0.253 | -0.17 | -11.24 | 36.22 | -0.31 | 0.757 | -0.05 |
| Strength Negative FC | 1.71 | 4.84 | 0.35 | 0.725 | 0.07 | -447.67 | 359.42 | -1.25 | 0.218 | -0.27 |

Note: Results for γ = 1.0 reported in main text

**Table S8. Correlations with PSAT and GPA for Modularity Difference Scores**

| | PSAT | GPA |
|---|---|---|
| **$Q_{Reasoning}$ - $Q_{Rest}$** | | |
| Q γ=0.5 | 0.311 | 0.380 |
| | 0.022 | 0.004 |
| Q γ=1.0 | 0.318 | 0.463 |
| | 0.019 | 0.000 |
| Q γ=1.5 | 0.251 | 0.406 |
| | 0.068 | 0.002 |
| Q γ=2.0 | 0.258 | 0.429 |
| | 0.060 | 0.001 |
| **$Q_{MRT}$ - $Q_{Rest}$** | | |
| Q γ=0.5 | 0.272 | 0.267 |
| | 0.032 | 0.033 |
| Q γ=1.0 | 0.190 | 0.271 |
| | 0.138 | 0.030 |
| Q γ=1.5 | 0.150 | 0.264 |
| | 0.246 | 0.035 |
| Q γ=2.0 | 0.181 | 0.339 |
| | 0.160 | 0.006 |

**Note**: *r* value indicated on top row, *p* value indicated on bottom row. Results very similar to those from task-based Q alone.
